# Tetrapod species-area relationships across the Cretaceous-Paleogene mass extinction

**DOI:** 10.1101/2024.09.13.612886

**Authors:** Roger Adam Close, Bouwe Rutger Reijenga

**Author notes:** R.A.C. and B.R.R designed research; R.A.C. and B.R.R performed research; R.A.C. and B.R.R analyzed data; R.A.C. and B.R.R and wrote the paper. The authors declare no competing interest.

## Abstract

Mass extinctions are rare but catastrophic events that profoundly disrupt biodiversity. Widely-accepted consequences of mass extinctions, such as biodiversity loss and the appearance of temporary ‘disaster taxa,’ imply that nested species-area relationships (SARs, or how biodiversity scales with area) should change dramatically across these events: specifically, both the slope (reflecting the rate of accumulation of new species with increasing area) and intercept (reflecting the density of species at local scales) of the power-law relationship should decrease. However, these hypotheses have not been tested, and the contribution of variation in the SAR to diversity dynamics in deep time has been neglected. We use fossil data to quantify nested SARs in North American terrestrial tetrapods through the Cretaceous-Paleogene (K/Pg) mass extinction (Campanian–Ypresian). SARs vary substantially through time and among groups. In the pre-extinction interval (Maastrichtian), unusually shallow SAR slopes (indicating low beta diversity or provinciality) drive low total regional diversity in dinosaurs, mammals and other tetrapods. In the immediate post-extinction interval (Danian), the explosive diversification of mammals drove high regional diversity via a large increase in SAR slope (indicating higher beta diversity or provinciality), and only a limited increase in SAR intercept (suggesting limited diversity change at small scales). This contradicts the expectation that post-extinction biotas should be regionally homogenized by the spread of disaster taxa and impoverished by diversity loss. This early post-extinction increase in SAR slope was followed in the Thanetian–Selandian (*∼*4.4. myr later) by increases in the intercept, indicating that diversity dynamics at local and regional scales did not change in synchrony. These results demonstrate the importance of SARs for understanding deep-time diversity dynamics, particularly the spatial dynamics of recovery from mass extinctions.

---

The Cretaceous–Paleogene mass extinction (K/Pg) was one of the most influential events in the history of Phanerozoic biodiversity. On land, this event caused the demise of several major groups, including all non-avian dinosaurs (1–5). Leading hypotheses of the causes of the K/Pg mass extinction include a large bolide impact in the Yucatan peninsula, which resulted in abrupt largescale environmental deterioration (2, 3), and the eruption of *>*1.1 million km^3^ of continental flood basalts over a period of *∼*750,000 years in the Deccan volcanic province (4, 6). Counter-intuitively, and perhaps uniquely among mass extinction events, the K/Pg event catalyzed rapid net increases in biodiversity, over timescales of hundreds of thousands to millions of years. The explosive radiation of mammals (7–11), in particular, resulted in a stepwise increase in the diversity of terrestrial (=non-flying, non-marine) vertebrates, establishing a new diversity equilibrium on land that was maintained throughout the Cenozoic (7, 12–15). However, we do not yet understand whether this post-extinction diversity rebound was driven solely by changes in local richness, or by changes to the spatial scaling of diversity.

Past studies of terrestrial tetrapod diversity across the K/Pg have investigated patterns at a variety of discrete spatial scales, including at local (15), regional and continental (12–14), and global (16, 17) levels. However, diversity increases with the area over which it is measured [e.g., (18)]. This ubiquitous pattern is formalized as the nested species-area relationship [SAR; (19)] which, over intermediate spatial scales, is commonly characterized using a power-law model (18). The form of the nested SAR reveals rich information about the spatial structure of diversity in continental settings, and especially about the links between local- and regional-scale diversity. Variation in regional-scale (gamma) diversity is an emergent phenomenon, driven by some combination of (1) changes in richness at small spatial scales (e.g., the richness of local communities, or alpha diversity), which causes the intercept of the nested SAR to vary; and (2) changes to the rate at which diversity scales with area, which causes the slope of the nested SAR to vary. The rate at which diversity scales with area in continental settings is a function of the level of spatial homogeneity of species distributions. The geographic turnover of species’ identities is often termed ‘beta diversity’ or, at larger scales, provinciality, and can be quantified directly for pairs of samples using a range of indices (20). The slope of the power-law relationship in nested SARs is an effective way to summarize information about species’ distributions across spatial scales (19). Understanding how diversity scales with area is especially important in the fossil record — species-area relationships are ubiquitous both today and in the past. Therefore, apparent changes in regional-scale diversity may be artifacts of variation in the geographic distribution of sampled fossil localities through time, which is known to be substantial (13–15, 21). For this reason, it is important to explicitly quantify spatial structure when estimating biodiversity dynamics through deep time.

Mass extinctions profoundly impact biodiversity at all spatial scales (22–25). By disrupting and subsequently restructuring ecological communities (22–26), and by upending existing patterns of species distributions (27, 28), these events have the potential to substantially alter key macroecological phenomena, including species-area relationships. For example, the selective extinction of species with smaller geographic ranges (25) would be expected to result in flatter SAR slopes, by eliminating those species that make the greatest contribution to beta diversity. Furthermore, post-extinction faunas may be dominated by super-abundant and highly-cosmopolitan ‘bloom’ or ‘disaster’ taxa (26, 29, 30). If correct, this would be expected to result in greater spatial homogeneity of species distributions in post-extinction assemblages. Both of these effects should result in flatter SAR slopes, while the loss of species at local scales might be expected to lower SAR intercepts.

A bolide impact represents the most likely kill mechanism for the K/Pg extinction. However, longer-term environmental stressors may also have contributed to ecological disruption in the lead-up to this event (1, 31), including flood volcanism (4, 6), climate change (1, 32–35), and sea-level change (36–38). Consistent with this scenario, ecosystem modeling has suggested that Campanian–Maastrichtian dinosaur communities in North America were more vulnerable to disruption of primary productivity (31), and latest Cretaceous (Maastrichtian) dinosaur assemblages in North America exhibited low beta diversity (39).

However, despite considerable interest in the effects of mass extinctions on terrestrial communities and species’ distributions (26–28, 30, 40), and in the K/Pg in particular [e.g., refs (10, 41, 42)], no study has directly quantified the effects of mass extinction events on terrestrial species-area relationships. In fact, few studies have quantified how species-area relationships of any kind have varied through time [but see (43–45) for Miocene-Recent mammals]. To better understand the effect of mass extinctions on macroecological patterns and processes, we need a unified understanding of how biodiversity varies across a range of spatial scales.

Here, we use fossil occurrence data to analyze how species-area relationships for North American terrestrial tetrapods, and major subgroups, varied across a window of *∼*36 million years around the K/Pg boundary (Campanian–Ypresian; Fig. 1). This interval encompassed the lead-up to the K/Pg extinction, its aftermath and recovery. It witnessed substantial environmental change between the Campanian and Maastrichtian (1), including orogenic activity and sealevel change affecting the paleogeography of the Western Interior seaway, Deccan flood-volcanism (4), and long-term global cooling towards the end of the Cretaceous (1, 32–35); a period of dramatic climate fluctuations during the early Paleogene (46); and protracted ecosystem recovery (41) that was only complete by the early Eocene (47). We focus on the North American tetrapod record from this interval because it is intensively-studied, well-documented, and has consistently broad geographic scope. By quantifying variation in the form of the SAR, we are able to determine which spatial scales drove overall patterns of biodiversity change over this key interval, and variation among major taxonomic groups, including non-avian dinosaurs, mammals and other taxa.

**Fig. 1.**
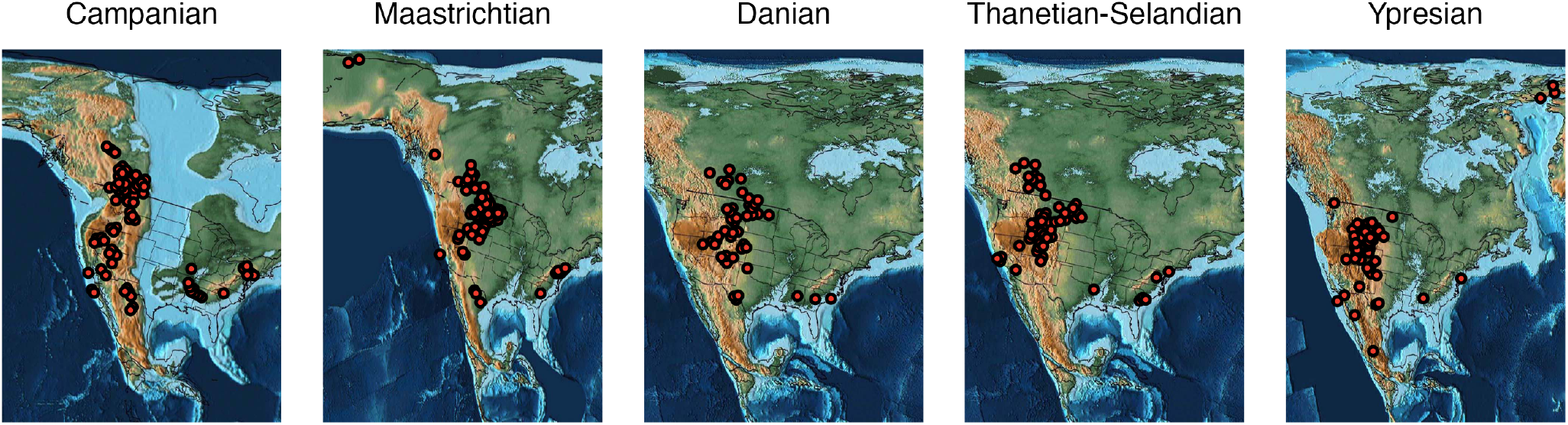
Variation in fossil sampling through time from the Campanian to the Ypresian. Red points represent coordinates of the mid-points of equal-area grid cells where terrestrial tetrapod fossils have been recovered. Paleogeographic maps were obtained from the PALEOMAP PaleoAtlas (48, 49) and cropped to the sampling extent of the fossil record.

## Results

Our results show a substantial shift in the species-area relationship (SAR) for terrestrial tetrapods over the K/Pg boundary. The slope of the SAR increased abruptly during the earliest Cenozoic (Danian; Figs 2–3). This indicates that species were much less homogeneously distributed in the Danian than they were during the latest Cretaceous (Maastrichtian), being instead more similar to that seen in the preceding Campanian interval. Tetrapods as a whole show little change in intercept in the Danian, but in mammals this increase in slope is accompanied by a near twofold increase in intercept (Figs 2–3) [consistent with ref (15)]. This jointly explains the large increase in regional-scale diversity for terrestrial tetrapods in the earliest Cenozoic. This increase in diversity was driven by the post-extinction radiation of mammals, which ultimately experienced a fourfold rise in the intercept of their SAR between the Maastrichtian and the Ypresian (Figs 2–3), indicating a quadrupling of mammal diversity within local communities into the Eocene. Patterns in minor groups (clades with lower diversity or less well-sampled fossil records) are less clear-cut (Fig. S1), either because they are characteristically species-poor (crocodilians, turtles) or because their fossil records are less well sampled (squamates, lissamphibians). Because of their data limitations, we focus primarily on the interpretation of the major groups and aggregated groups. However, it is clear that none of these groups experienced the rapid and emphatic post-extinction rebound seen in mammals.

**Fig. 2.**
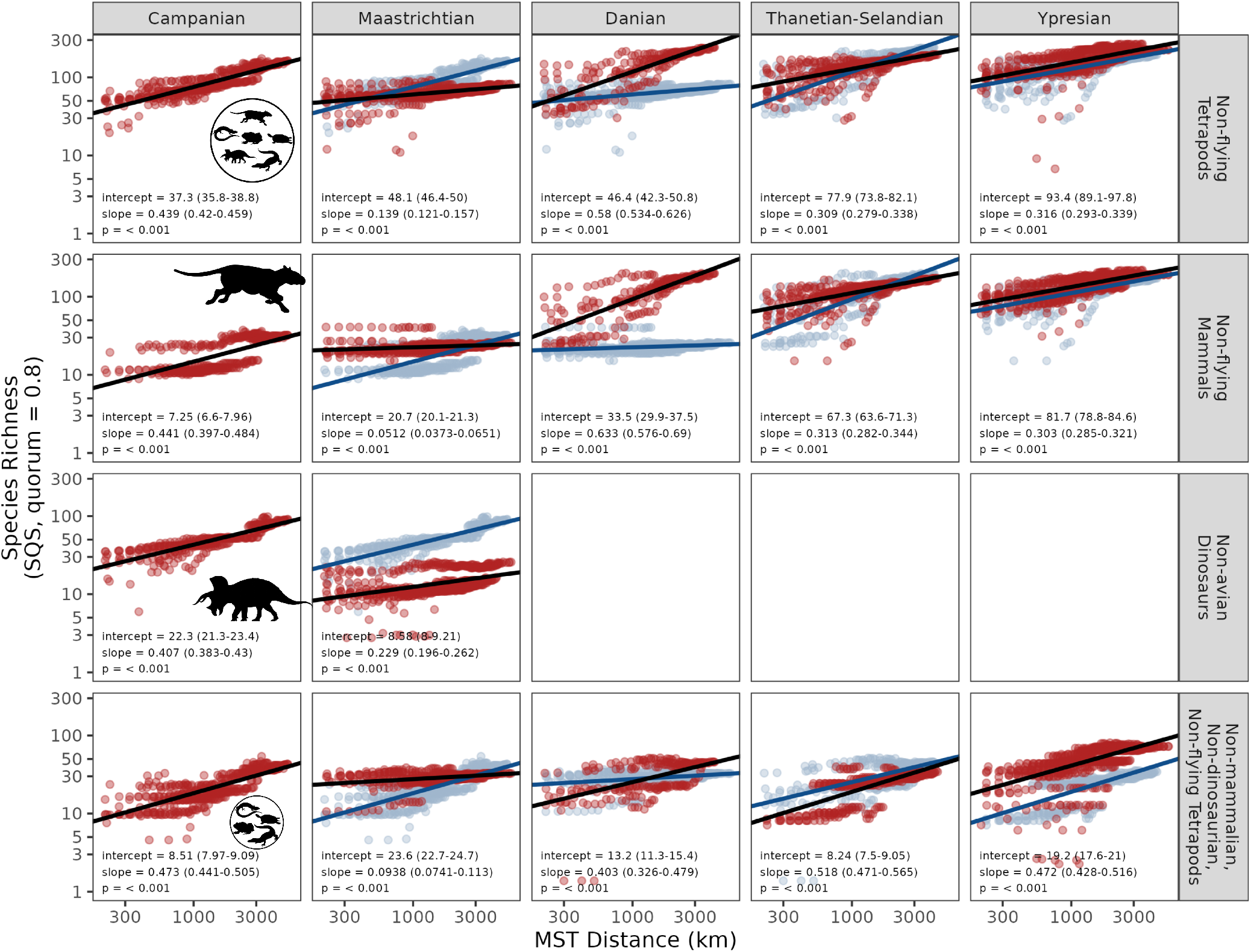
Species-area relationships for North American terrestrial tetrapods and major subgroups across the Late Cretaceous–early Paleogene. Columns represent the composite time bins, and rows represent the taxonomic groups. Each facet shows the nested species-area relationship for the respective group and bin (red), and the previous bin if relevant (blue). Ordinary least-squares regression fits and estimated parameters are shown for the current (black) and previous bin (blue). Species richness estimates were obtained using SQS at a quorum of 0.8

**Fig. 3.**
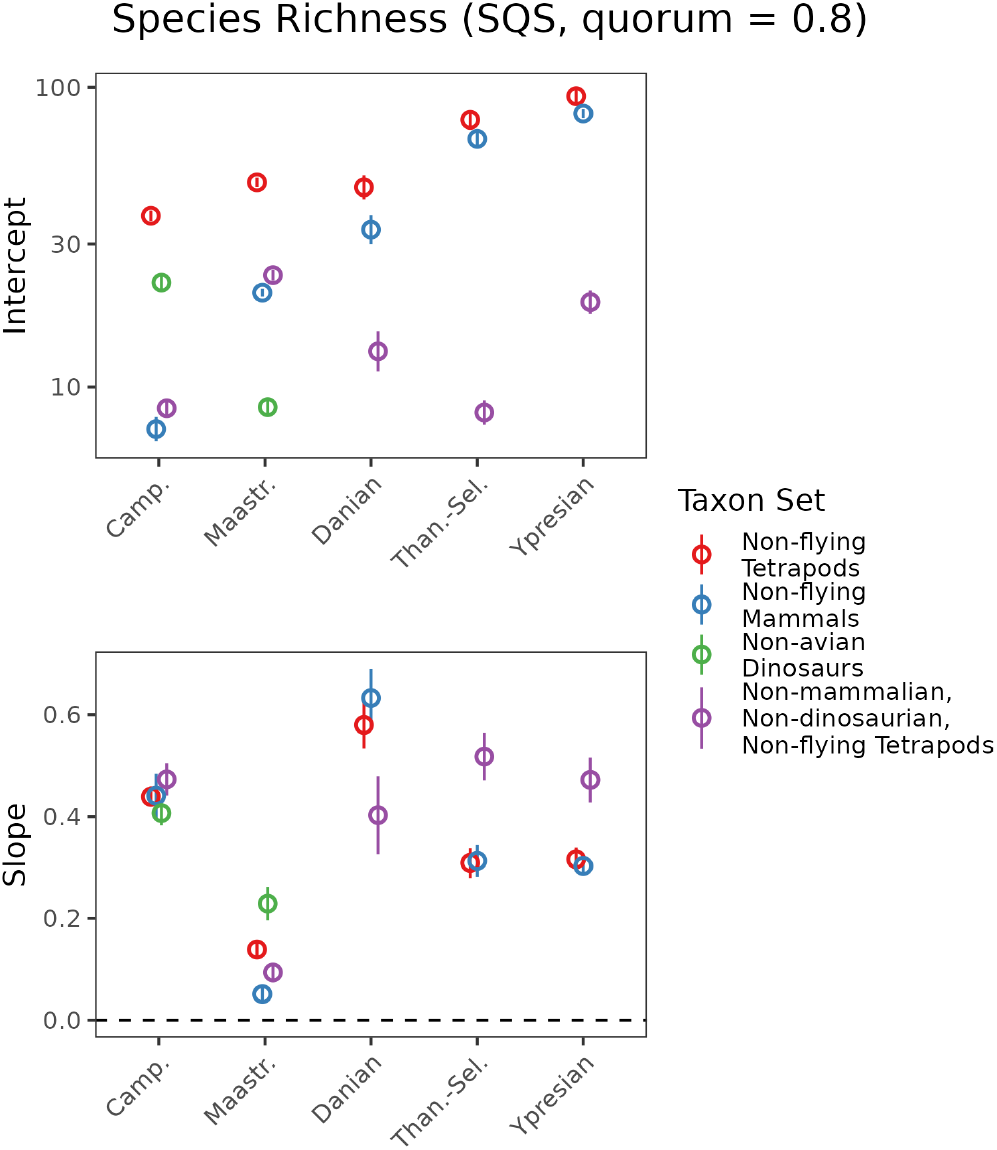
Slope and intercept parameters for species-area relationships (richness estimates obtained using SQS at a quorum of 0.8) for North American terrestrial tetrapods and major subgroups across the Late Cretaceous–early Paleogene.

We also find evidence for a comparatively short-lived departure towards a substantially shallower slope during the latest Cretaceous (Maastrichtian), indicating a substantial decrease in beta diversity or provinciality before the K/Pg boundary (Figs 2–3 and S1; statistical tests of significance in slope between successive pairs of time bins are given in Table 1). Slopes for SARs of terrestrial tetrapods as a whole (Figs 2–3 and S1) decreased from *∼*0.439 in the Campanian to 0.139 in the Maastrichtian, and 95% confidence intervals do not overlap. SAR slopes were shallower during the Maastrichtian in nearly every major tetrapod group, including dinosaurs, lissamphibians, mammals, squamates and turtles, and aggregated minor groups (non-mammalian, non-dinosaurian, non-flying tetrapods; Fig S1). The shallow SAR slope in the Maastrichtian (suggesting lower beta diversity or decreased provinciality) produces the lowest regional-scale diversity of all the intervals examined here—for terrestrial tetrapods, dinosaurs, squamates, and to a lesser degree, mammals. Intercepts of SARs show more variation among groups. In tetrapods as a whole, the intercept increases slightly from the Campanian to the Maastrichtian, although diversity estimates for individual subsampled spatial regions broadly overlap between the two intervals. Dinosaurs and mammals, however, show opposing patterns, with a modest increase in intercept for mammals and a large decrease in dinosaurs.

**Table 1.**
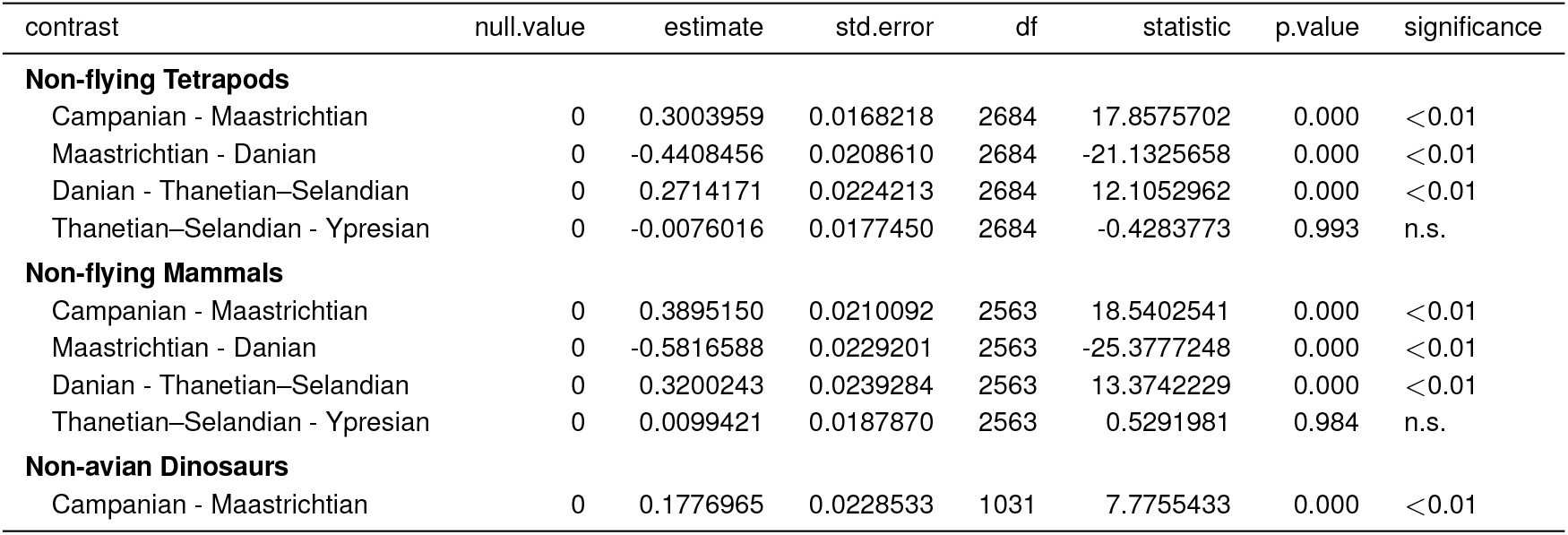
OLS slope changes between time bins.

Although the SAR slope did increase in the Danian, following the K/Pg, longer-term increases in regional species richness between the Mesozoic and Cenozoic result more strongly from increases in intercept rather than slope. Although large and statistically-significant differences are recovered between slopes for the Maastrichtian and Danian, slopes are very similar between the Campanian and either the Thanetian–Selandian or Ypresian. Furthermore, Early Paleocene (Danian) increases in the slope of SARs were followed by an increase in the intercept and a sharp relaxation of the slope to lower levels later in the Paleocene (Thanetian– Selandian). Subsequent changes in slope and intercept for tetrapods and mammals between the Thanetian–Selandian and Ypresian are relatively minor. While the slope decreases significantly from the Danian to the Thanetian–Selandian, from the Thanetian–Selandian to the Ypresian the slope in tetrapods and mammals remains stable. On the other hand, intercepts increase modestly, resulting in slightly higher gamma diversity at the largest comparable spatial scales.

### Discussion

We present the first direct reconstructions of species-area relationships (SARs) across a major extinction event, using an approach for quantifying nested species-area relationships that extracts rich information about the spatial scaling of diversity in the fossil record. Furthermore, by comparing empirical SARs with geographically-scrambled null distributions and Spatially-Explicit Neutral Models (SENMs), we show that SARs generated using face-value (=‘raw’ or ‘unstandardized’) species counts are misleading: due to undersampling, they predominantly represent sample-accumulation curves, and cannot be interpreted as real biological species-area relationships. In contrast, SAR relationships estimated here using sampling-standardised richness estimates (SQS) do not suffer from this problem (Fig. S2).

### A. Decrease in species-area slope in the Maastrichtian

Variation in the slope of the terrestrial tetrapod SAR through the latest Cretaceous–early Cenozoic suggests an important role for variation in the spatial scaling of diversity in the dynamics of mass extinction and recovery. An especially shallow slope occurs in the latest Cretaceous (Maastrichtian), implying low beta diversity during the lead-up to the mass extinction. This contrasts with substantially steeper slopes (implying higher beta diversity) during the Campanian and earliest Cenozoic (Danian), an effect that is observed across all taxonomic groupings (Fig. 2, Fig. S1). In terrestrial tetrapods as a whole, the shallower slope primarily reflects diminished regional-scale diversity relative to the Campanian (Fig. 2), while in mammals and other non-mammalian, non-dinosaurian tetrapods, it reflects only a modest decrease in regional-scale richness, coupled with a large increase in richness at small spatial scales (i.e., a higher intercept; Fig. S1).

Dinosaurs show a pronounced decrease in both the slope and intercept of their SAR in the Maastrichtian. This is a feature that cannot easily be discerned by estimating diversity only for narrow-ranging, discrete spatial scales (Fig. S1). This finding is consistent with a previously-documented decrease from the Campanian to the Maastrichtian in the local (15) and regional-scale (14) richness of North American dinosaurs. Vavrek and Larson (39) also suggested that beta diversity in Maastrichtian dinosaurs of western North America was low. However, this inference was made by comparing Maastrichtian beta diversity to extant North American birds, and without reference to other fossil data. Here, by contrast, we show that beta diversity (as measured by the SAR slope) in dinosaurs decreased from the Campanian into the Maastrichtian, and that other tetrapod groups experienced similar decreases over this interval.

This trend may capture a true biological signal of decreasing beta diversity, perhaps driven by tectonic (34) or climatic changes (4) from the preceding Campanian. Alternatively, the trend may reflect differences in the spatial, environmental and temporal structure of the Campanian and Maastrichtian fossil records of North America (50, 51). By explicitly quantifying nested species-area relationships, we address issues arising from differences in spatial coverage between stages, but future analyses could combine this spatially-explicit approach with information on environmental and depositional settings and taphonomic conditions.

### B. Patterns across the K/Pg boundary

We show that the end-Cretaceous mass extinction catalyzed increases in both the intercept and slope of the SAR in North American terrestrial tetrapods at the temporal resolution of our study (Fig. 2). The SAR slope underwent a large, early increase between the Maastrichtian and the Danian, followed by an increase in the SAR intercept, and relaxation of the slope, up to 4.4 million years later (between the Danian and Thanetian– Selandian; Fig. 2). This temporal decoupling of SAR slope and intercept indicates a substantial contribution of variation in beta diversity of the dynamics of regional diversity change in North America during the K/Pg mass extinction and recovery.

Although it is likely that we can reliably estimate SAR slopes and intercepts, as shown by our SENM simulations (Figs S3, S4, S5, and S6), the temporal resolution of our data might influence our inferences. Specifically, consider the increased SAR slope of the Danian. If rapid temporal turnover of species occurred during this geological stage, this might induce heightened spatial turnover if different localities were sampled from different time points within the stage. We do not have any prior expectation that this would particularly impact the Danian, or how this could systematically bias our results. However, future work would benefit from more finely-resolved temporal resolution, although likely at the cost of spatial resolution.

The increase in intercept and slope between the Maastrichtian and the Danian reflects modestly higher tetrapod diversity at small (*∼*100 km MST distance) spatial scales, coupled with more substantial increases at large (*>*1000 kmMST distance) spatial scales. These changes are driven by a pronounced increase in the mammalian SAR intercept. Other tetrapod groups examined here did not experience a rapid post-extinction rebound (Fig. 2, Fig. S1), consistent with previous findings for groups such as squamates (52) and non-marine crocodylians (53). Changes to the form of the SAR through the Cenozoic in tetrapods and mammals are relatively minor.

Our findings are consistent with recent work on tetrapod species richness at local to regional spatial scales, which documented twofold to fourfold increases in terrestrial tetrapod diversity in the earliest Cenozoic, driven by the explosive radiation of mammals (9, 12–15, 41, 54). Together, this work suggests that diversity equilibria for terrestrial tetrapods were reset at local and/or regional spatial scales. A similar phase-shift towards higher diversity across the K/Pg was recently documented for marine animals at regional spatial scales, driven by the explosive radiation of gastropods (21). Taken together, these results contribute to a revised understanding of the macroevolutionary role of the end-Cretaceous extinction. The emerging picture contradicts classical expectations that mass extinctions primarily eliminate biodiversity, and instead suggests that the K/Pg mass extinction was a major generative force, ultimately creating higher levels of biodiversity—in large part due to exceptional radiations by a small number of taxonomic groups—that persisted through the Cenozoic.

## Materials and Methods

### Data

Fossil occurrence data for terrestrial tetrapods (i.e., nonflying, non-marine groups) were downloaded from the Paleobiology Database (http://www.paleobiodb.org/) on 17 November 2023. All procedures for downloading and processing occurrence data (e.g., to excludeunsuitable records) are identical to those described in (14), except that the analysis was restricted to North America over the interval spanning the Campanian to the Ypresian. As part of this procedure, the occurrence records were parsed into sets for major tetrapod subgroups (‘dinosaurs’ = Dinosauria excluding Aves; ‘mammals’ = Mammaliamorpha excluding Chiroptera; ‘squamates’= Squamata, ‘crocodilians’ = Crocodylomorpha; ‘turtles’ = Testudinata; ‘lissamphibians’ = Lissamphibia). Flying taxa (Aves, Chiroptera and Pterosauria) were excluded because their fossil record is inadequate for most intervals and regions, and dominated by Lagerstätten (13–15). We take the lack of information about flying taxa into account when interpreting our results. The analysis was conducted at species-level. The final cleaned occurrence dataset comprised 14,887 occurrences from 3,766 collections, representing 1,661 species. Occurrence data were binned into nominally equal-length composite time bins, created by combining stratigraphic stages, following (14). Over our study interval, however, only the Thanetian–Selandian bin comprises more than a single stratigraphic stage.

### Spatial subsampling

We reconstructed nested SARs by estimating diversity within spatial regions of known paleogeographic extent, as calculated from the paleocoordinates of fossil localities. Each subsampled spatial region consists of a set of geographically-adjacent fossil localities. Following our previous work on spatially-explicit diversity estimation (13, 14, 21), we used minimum spanning tree (MST) distance in kilometers as our measure of spatial extent [see (13)]. Subsampled spatial regions were constructed using an algorithm [modified from (55)] that identifies all unique nested sets of directly adjacent fossil localities. To reduce computational complexity, paleocoordinates for fossil localities were binned into 100 km equal-size hexagonal/pentagonal grid cells [using the R package dggridR (56)] prior to spatial subsampling. Midpoints of these 100 km grid cells were used to define the spatial points used in our spatial subsampling algorithm. Our algorithm comprises the following steps: (1) first, a spatial point is randomly chosen as a starting location; (2) the nearest spatial point is then identified (chosen at random if there are two or more equidistant points), and these two points are used to define the first subsampled spatial region; (3) the nearest spatial point to the first two points is identified, and this set of three points is used to define the second subsampled spatial region; and (4) this procedure continues, point by point, until all spatial points have been added. This procedure is then repeated using every possible starting location, and any duplicate paleogeographic regions are removed. Our implementation of this algorithm is identical to that described in (14) and (21), and full details are given there. Note that we did not use the subsequent region-clustering steps employed by (14) and (21), as these are only suitable for analyzing regional-scale diversity at discrete spatial scales. To avoid estimating diversity from regions containing widely-separated clusters of localities, we screened our regions to ensure that the maximum nearest-neighbor distances between localities was less than 1,000 (see Supplementary Materials: Methods). Using this spatial subsampling procedure, we identified a total of 2,792 distinct regions between *∼*150 km and *∼*5,500 km MST distance across all time intervals.

### Diversity estimation

Species richness was estimated using Shareholder Quorum Subsampling [SQS; (57–61)] for each subsampled spatial region, using the R package iNEXT (62). We present results using a quorum of 0.8 in the main text. This balances richness estimator performance with data availability. For comparative purposes, face-value counts of species (=‘raw’ or uncorrected species richness) were also calculated for each subsampled spatial region. However, simulated null distributions suggest that the SARs using these face-value richness estimates reflect sample-accumulation curves to a greater extent than they do biological species-area relationships (Supplementary Materials: Results), so we focus on results obtained using SQS.

### Estimating empirical species-area relationships

SARs were constructed by plotting diversity estimates for each subsampled spatial region as a function of paleogeographic extent (km MST distance) with log-log axes. The form of the SAR was characterised using ordinary least-squares (OLS) regression, as per (55). The slope of the relationship corresponds to the *z* parameter in the classic power-law equation describing linear SARs, and the intercept to the *c* parameter (18, 63).

We focus our interpretation on the intercept of the OLS regression line at an MST distance of 100 km, which corresponds to the minimum size of the subsampled spatial regions we analyzed. For convenience, we refer to this as the ‘intercept’, and use it as an estimate of local diversity at a scale of approximately 100 km. Extrapolating relationships to smaller scales can produce counter-intuitive results, because increases in the SAR slope can cause a decrease in the estimated intercept, even in the absence of any decrease in the species richness observed for the smallest subsampled spatial regions. Moreover, very small spatial scales lie outside the Arrhenius Zone (the range of spatial scales in nested SARs where linear changes in diversity as a function of area are observed in log-log space (18, 19, 63)), and at smaller scales, a nonlinear relationship is observed (19). For this reason, the intercept of the SAR should here be interpreted as diversity at the smallest measured spatial scales, rather than local richness.

We tested for differences in slopes of successive bins using an interaction with covariates analysis implemented via the function ‘emtrends()’ in the R package ‘emmeans’.

The slope of the power-law relationship in a nested SAR can be straightforwardly interpreted as the average rate of scaling of diversity with area and of average beta diversity across the spatial scales being analyzed. However, the intercept of the nested SAR cannot be solely interpreted in terms of changes in local richness. There are two reasons for this. Firstly, in this study, the smallest spatial scales quantified (*∼*100 km summed MST distance) are larger than those needed to quantify local richness. As a result, beta diversity at very small scales (e.g., among adjacent habitats) could drive changes in the intercept even if local richness *per se* did not change. Secondly, changes to the slope can also affect the estimated intercept at zero kilometers, even if richness at the smallest measured spatial scales did not change. For example, increasing the slope will also decrease the intercept, while diminishing the slope will increase the intercept. For these reasons, we do not use the traditional estimate of the intercept (i.e., the slope of the regression line at zero), and instead use the estimate at the smallest measured spatial scale of 100 km.

### Null distributions

Incomplete sampling of diversity introduces a signal of sample-accumulation to nested SARs, which is independent of the true relationship between diversity and area. This results from increasing sample coverage at larger spatial scales, and may exaggerate slopes estimated from empirical data, especially when using face-value counts of species (=raw or uncorrected species richness). To assess the potential impacts of this, we compared our empirical SARs to a null distribution generated by randomizing the spatial structure of the fossil record. This was achieved by randomly reassigning the palaeocoordinates of collections within each bin, without replacement, to remove the effects of spatial structure and beta diversity. This allowed comparison of our empirical regression relationships to those based on null distributions in which no SAR should exist, except due to the effects of sample incompleteness. These null distributions show that SARs estimated using SQS contain a much weaker sample-accumulation signal than those using face-value richness (Supplementary Materials: Results; Fig S2), and we therefore focus exclusively on SQS when interpreting results. Furthermore, null distributions for minor groups, even when aggregated (non-mammalian, non-dinosaurian, non-flying tetrapods), suggest that SAR parameter estimates for some intervals may be less reliable than those for the major groups (e.g. Thanetian-Selandian, Fig. S2). For this reason, we focus our interpretation more on the major groups (non-flying tetrapods, mammals, and dinosaurs).

### Spatially Explicit Neutral Models

The fossil record shows variation through time in the number of localities sampled, their geographic dispersion, and the intensity with which they are sampled. Our ability to infer changes in SARs through geological time may be impacted if sampling is not sufficiently extensive or complete. We used spatially explicit neutral models [SENMs; (64, 65)] to assess if the empirical sampling structure is sufficient to infer the shape of and changes in SARs through time. SENMs allow for the efficient generation of assemblages of many individuals, from small islands to continents, and have been shown to generate realistic SARs (64, 66, 67). Assemblages simulated across the entirety of the North American subcontinent could then be subsampled to reflect the sampling extent and intensity of the fossil record, so that we knew both the true and subsampled SAR. Our goals were to investigate if there was (1) any inherent bias in estimation of SAR slope and intercept caused by variation in sampling through time, (2) if local-to-global SARs could be accurately inferred from regional fossil SARs, (3) if excluding clusters of localities with a nearest-neighbor distance greater than 1,000 km was justified, and (4) to further investigate if face-value counts of species or those estimated using SQS were more accurate.

We briefly summarise the results of these analyses here. (1) We found that face-value counts lead to misleading estimates of both slopes and intercepts, showing sudden increases or decreases when the actual slopes and intercepts show no trend (Figs S3, S4, S5, and S6). (2) We also found that if slopes and intercepts are estimated from only the geographic cells that contain fossils, the estimates do not show an apparent bias (Figs S3, S4, S5, and S6). (3) When we downsample these cells to reflect the sampling heterogeneity through space (e.g. variation in the number of collections and occurrences), and correct for this unevenness with SQS, we find that both slope and intercept are underestimated systematically. However, no apparent bias arises in the estimates. The Danian shows greater variability then any of the other time bins. The estimates become more precise when dispersal distance increases (Figs S3, S4, S5, and S6). (4) Including long branches (i.e., grouping clusters of cells located further than 1,000km from any other cells) causes estimations to poorly reflect the slopes and intercepts of SARs against which they are compared (as they cover a larger area for which there is limited information; Figs S3, S4, S5, and S6). (5) Lastly, a comparison between local-to-continental SARs and the regional SARs estimated from the fossil sites (completely sampled), shows that although the regional SAR shifts with changing parameter values (reflecting the continental trends), the regional SAR has a limited ability to pick up all continental trends, often overestimating the intercept and underestimating the slope, as species poorer areas are not captured. (*SI Appendix*, Figs S7, S8, S9, and S10)

## ACKNOWLEDGMENTS

We thank J Rosindell and S Thompson for help with Rcoalescence, and everyone that has contributed to the Paleobiology database. this is PBDB publication number

## Notes

### Competing Interest Statement

The authors have declared no competing interest.

