## Supplementary Information for "Tetrapod species-area relationships across the Cretaceous-Paleogene mass extinction"

**This PDF file includes:**

Supporting text
Figs. S1 to S15
SI References

### Supporting Information Text

### 1. Supplementary Methods

**A. Identifying contiguous regions for estimating species-area relationships.** Quantifying SARs for sets of fossil localities which span multiple distinct spatial aggregations (i.e., in which groupings of localities are separated by longer distances devoid of sampling) is undesirable, because multiple distinct relationships are more likely to be superimposed. To avoid this, we identified distinct spatial aggregations of fossil localities using minimum-spanning trees (MSTs). For each interval, a global MST was calculated using the paleocoordinates of the fossil localities (in this case, the 'global' MST only encompasses North America; Fig. 1). The global MST was then split into subtrees by removing branches longer than 1,000 km. Subtrees with fewer than 10 MST nodes (i.e., binned paleocoordinates from which fossil localities are known) were excluded from the analysis due to inadequate data. In our case (North America from the Campanian to the Ypresian), this resulted in just one SAR per equal-length bin. Only subsampled spatial regions that were entirely contained within the identified subtrees were used to construct SARs.

**B. Spatially Explicit Neutral Models.** SENMs are a powerful but underused tool to simulate the distributions of biodiversity at the level of individuals (1), and are used here to investigate the effects of spatial sampling structure on the inferred shape of species-area relationships in the fossil record. In contrast to more complex ecological individual-based models, they benefit from coalescence methods that improve computational tractability (2). The key assumption of neutral models that makes them efficient is that individuals are ecologically equivalent regardless of species identity (i.e., all species are demographically identical in terms of their per-capita birth, death, dispersal and speciation rates) (3, 4). Consequentially, biodiversity is structured by a combination of speciation, dispersal and fluctuation in abundances of species. Despite neutral theory explicitly ignoring ecological niche differences between species, SENMs have been shown to result in realistic SARs (5–7). However, the limitation of this approach is that if the world is not neutral, the processes that cause changes in slopes and intercepts in neutral simulations might not be the same processes that cause variation in empirical SARs.

The simulation loop under SENMs is detailed elsewhere (8) but the mechanics are summarized here: An individual perishes, creating empty space for a new individual to be born. The species identity of this newborn individual is derived from neighboring individuals for which the likelihood decreases by the geographical distances between parent and offspring. Occasionally, with probability $\nu$, speciation occurs, and the offspring is assigned a distinct species identity from its parent. Over time, this results in individuals that are geographically distant to have a higher probability of having become different species. Entire species communities may therefore also show turnover in composition, strengthened by geographical barriers such as seaways that limit dispersal between communities even further.

We obtain information on geographical barriers and availability and connectivity of habitat cells from paleo-digital elevation models (paleoDEMS) at 80 to 50 Ma (9). PaleoDEMS describe the changing distribution of oceans, seas, lowlands and mountains, and provide an estimate where terrestrial tetrapods may have occurred over the K-Pg boundary. Habitat is constrained to any coordinates at a 0.1-degree resolution that are above sea level (> 0 m). For computational tractability, and having data in line with the empirical record, we bin these coordinates in equal-area grid cells spaced 100 km apart (**?** ). Only grid cells that are located on the North American subcontinent and contain at least one coordinate above sea level are used.

Our hexagonal grid cannot be accurately approximated by a square lattice where distances are measured in the number of cells between two points. We therefore use $n$ by $n$ dispersal matrices for each geological stage where $n$ equals the total number of habitat cells in each composite time bin. Row and column combinations highlight relative dispersal probabilities scaled between 0 and 1 between cells. The diagonal of the matrix is the probability to fill a gap in the cell that the individual is currently present in. This probability is higher if cells are more geographically isolated, as they are less likely to disperse to neighboring cells. The dispersal probabilities between cells are determined by a Gaussian dispersal kernel that describes dispersal probabilities based on the great-circle distance between cells. The width of this kernel determines how far individuals can disperse and can thus influence SAR shape directly.

SARs arising under SENMs can vary in shape and can reflect all three phases observed in empirical curves: an initial concave sampling phase at local scales, the Arrhenius power-law phase where diversity increases approximately linearly, and a convex phase at continental scales where regions separated by large distances contain non-overlapping biota (5, 6). In our simulations we aim for SARs that approximate linearity across our entire grid in absence of the Western Interior Seaway – the main dispersal barrier that may have had an influence on empirical SARs (10). We justify a linear expectation for simulated SARs as the area that we measure diversity over does not reflect the large intercontinental scales at which convex increases are expected, nor do they contain the small local scales required for concave increases (11). We therefore conducted preliminary exploration of parameter space to obtain SARs that reflect (*i*) linearity, (*ii*) scenarios of realistic total richness (i.e., around the same order of magnitude as contemporary richness of ∼2000 non-flying terrestrial tetrapod species), (*iii*) species abundance distributions where neither all species have the same abundance nor extreme imbalance, (*iv*) and plausible patterns of distance-decay where turnover in community composition with distance is neither completely non-existent (i.e. only cosmopolitans) nor complete at the shortest distances (Figs S11, S12, S13 and S14).

The two parameters that are varied in our SENMs to attain these relationships are speciation rate and dispersal distance. The per-capita speciation rate $\nu$ has a direct impact on total species richness, but as the total number of speciation events depends on the number of individuals, we keep the number of individuals per cell constant. Dispersal events happen with probability 1 - $\nu$, so is directly dependent on the frequency of speciation events. Speciation can create a high degree of endemicity due to newly arisen species consisting of a single individual, reducing average range sizes and skewing abundance distributions. Dispersal distance equally influences range sizes and abundances. Larger distances means that individuals can move further away. Initially when dispersal distance is increased, this increases range size as individuals of a species are not concentrated in a single community. However, greater dispersal distance also causes range fragmentation as the frequency of dispersal events remains constant. This also decreases the abundance of species in each cell and flattens abundance distributions. Although many shapes can be attained, SARs generally become more linear, and less convex/concave, when speciation rate increases and dispersal distance decreases (12). We vary speciation ($\nu$ = 1e-4, 2.5e-4, 5e-4, 7.5e-4, and 1e-3 at 100 individuals per cell), and distance ($\sigma$ = 25, 50, 100, and 150 km), and keep carrying capacity constant ($K$ = 500 individuals per cell) in the light of the four constraints mentioned above.

The realizations of our SENMs are descriptions of the count and frequency of species in each grid cell for the four paleoDEMs from 80 to 50 Ma. We calculate local-to-global SARs for these species inventories in the same fashion as described for the empirical data by, starting from each grid cell independently, cumulatively adding grid cells that are closest to the grid cell until all cells have been added to the cluster. We only keep clusters that have unique combinations of cells. We repeat this process for the regional SARs that respectively consist of (*i*) all grid cells where fossils have been sampled, (*ii*) grid cells in which the nearest-neighbor distance is no more than 1000 km, and (*iii*, *iv*) the respective convex hulls of the grid cells in (*i*) and (*ii*) containing all cells within the hull. OLS regressions were used to characterize the SAR slopes and intercepts on log-log axes. However, instead of using minimum spanning tree (MST) length, we correlate cluster richness to the summed path length (SPL). SPL is the cumulative minimum distance obtained from the clustering algorithm. We opt for using SPL instead of MST as SPL correlates strongly with MST and MST becomes computationally unfeasible when calculated for the entire continent (Fig S15). Note that for the regional SARs of the Danian and Thanetian–Selandian the same base simulation at 60 Ma is used, and only the spatial sampling changes. The Campanian, Maastrichtian and Ypresian respectively correspond to paleoDEMs at 80, 70 and 50 Ma.

The fossil record is highly heterogeneous in terms of the number of occurrences per grid cell or collection. Fossil occurrences are difficult to precisely translate to ecological presence-absences or frequency of a species in a community, because they reflect a complex combination of local richness, sampling effort, and fossil preservation. We approximate the sampling process of fossil occurrences in each grid cell by taking random draws from the individuals within each cell. The number of draws taken per grid cell is equal to the number of fossil collections, and the number of individuals sampled per draw is taken from a zero-truncated geometrical distribution with a shape equivalent to the number of occurrences per collection as found in the empirical record. We repeat this process 100 times for all parameter combinations and geological stages and calculate regional SARs based on face-value (i.e., raw) counts of species, and SQS set at a quorum of 0.8. Simulating variability in sampling across grid cells therefore gives us a clearer picture of how SARs are expected to vary.

### 2. Supplementary Results

**A. Null distributions.** Null distributions for SARs, in which spatial structure is destroyed by shuffling collections around paleocoordinates, show that relationships estimated using face-value taxon counts contain strong sample-accumulation curve component (Fig S2). Using face-value richness, the median slope of the null distribution is almost always comparable to the empirical slope, even though no relationship between richness and area should exist. In contrast, sampling-standardised SARs estimated using SQS do not suffer from this problem (Fig S2). Null distributions using SQS are, on average, almost perfectly flat, correctly inferring that no relationship between diversity and area exists. Conversely, the empirical relationships estimated using SQS are much steeper than their corresponding null distributions. We therefore focus our interpretation entirely on patterns estimated using SQS.

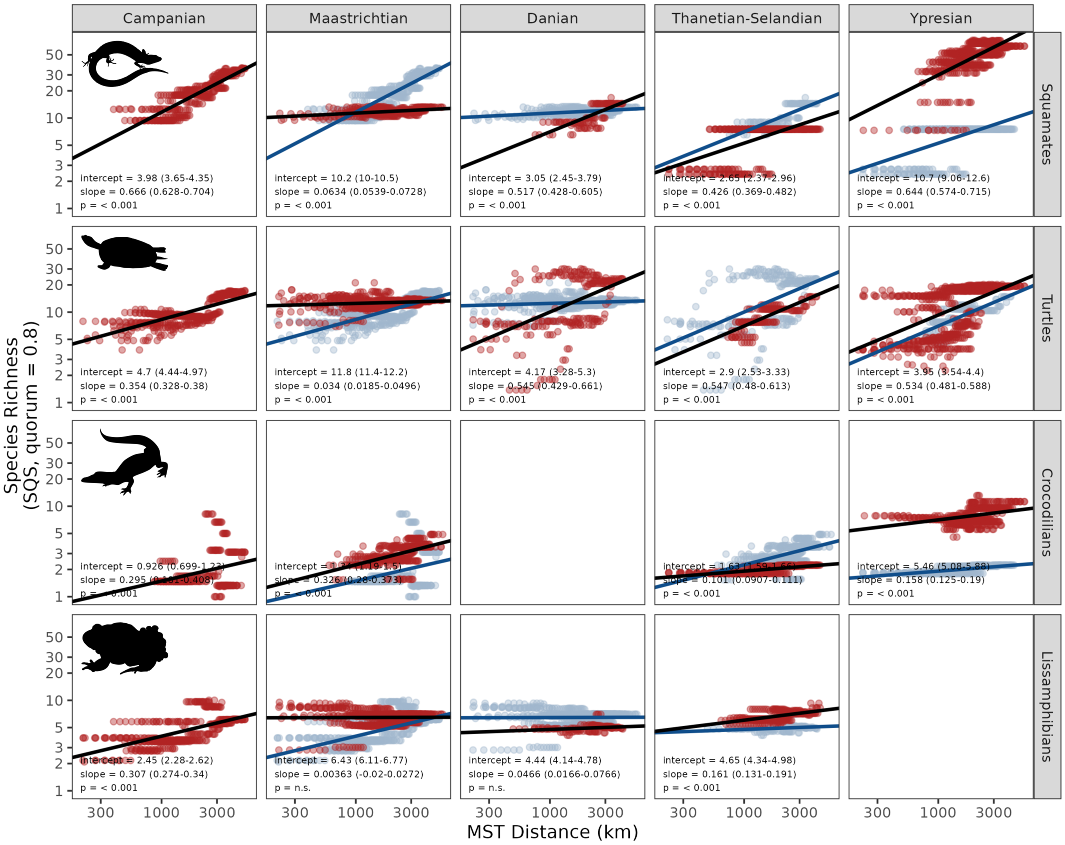


**Fig. S1.** Species-area relationships for less well-sampled subgroups (squamates, turtles, crocodilians, lissamphibians). Columns represent the composite time bins, and rows represent the taxonomic groups. Each facet shows the nested species-area relationship for the respective group and bin (red), and the previous bin if relevant (blue). Ordinary least-squares regression fits and estimated parameters are shown (black).

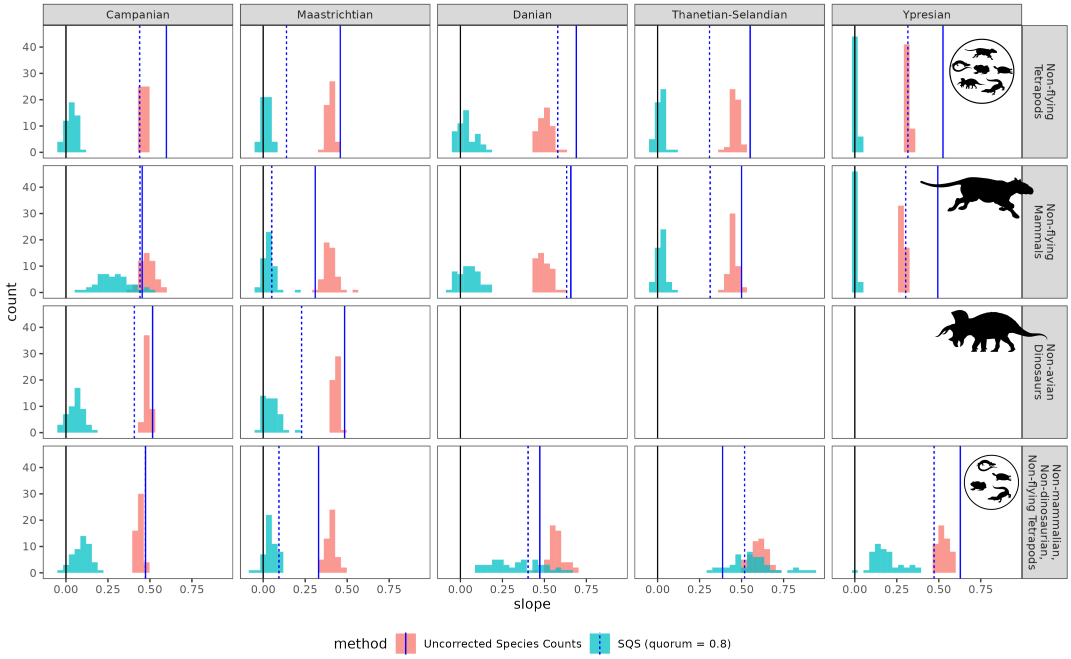


**Fig. S2.** Histograms showing distributions of slope estimates for the null models. Distributions are shown for slope estimates for uncorrected species counts (face-value diversity, red) and SQS (quorum = 0.8, blue). Dark blue lines show the empirical slope estimates for uncorrected (solid) and SQS (dashed) richness estimators. Black lines denote a slope of zero.

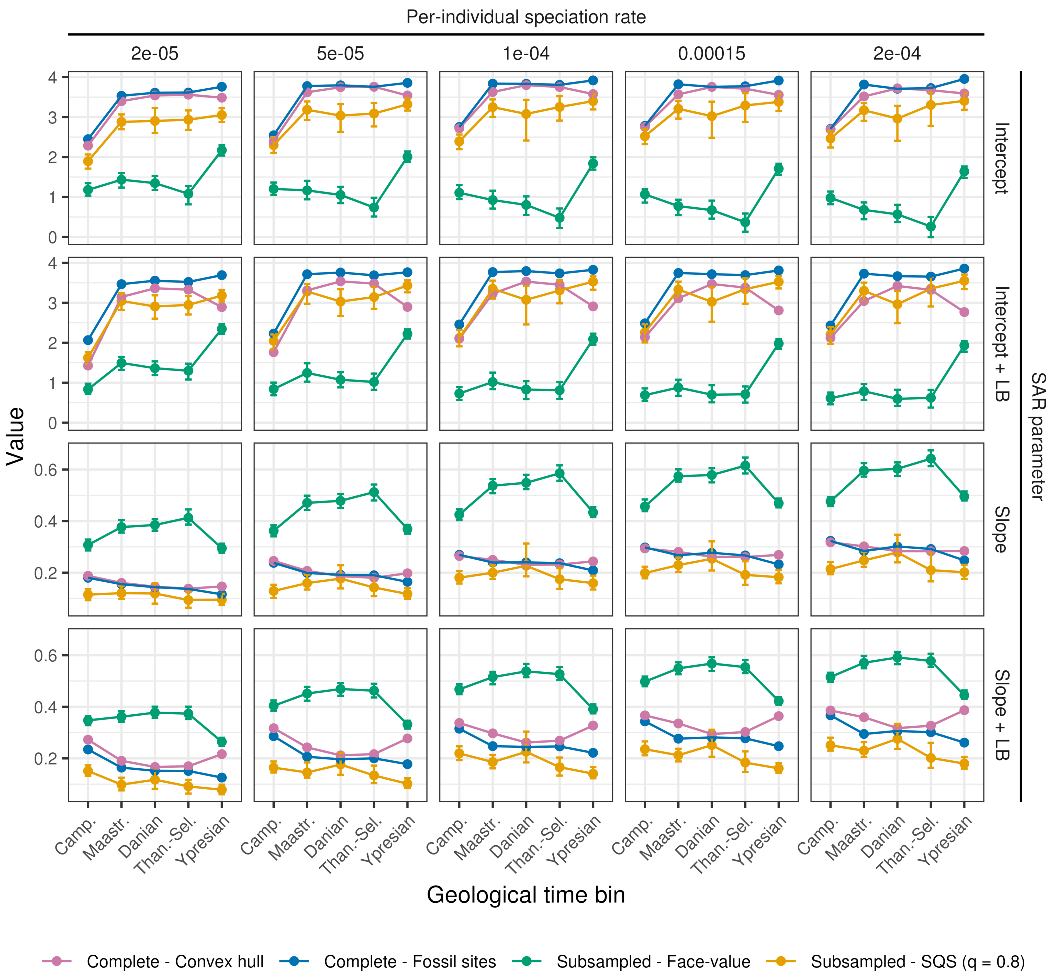


**Fig. S3.** Slopes and intercepts of SENMs for a dispersal distance $\sigma = 25$. OLS regression estimates are shown for SENM simulations that have been subsampled to the region of interest. Columns represent increasing per-individual speciation rates. Rows represent the intercepts and slope estimates of the regressions for the data sets with fossil sites at a great-circle distance of 1000km removed or included (+LB). Distinct colours respectively highlight SAR estimates for the full convex hull with all sites perfectly sampled (purple), complete sampling of all fossil sites (blue), sample sites are subsampled to reflect the number of collections and diversity is measured by face-value counts (green) or SQS (orange). Brackets show the 95% confidence intervals across 100 randomizations.

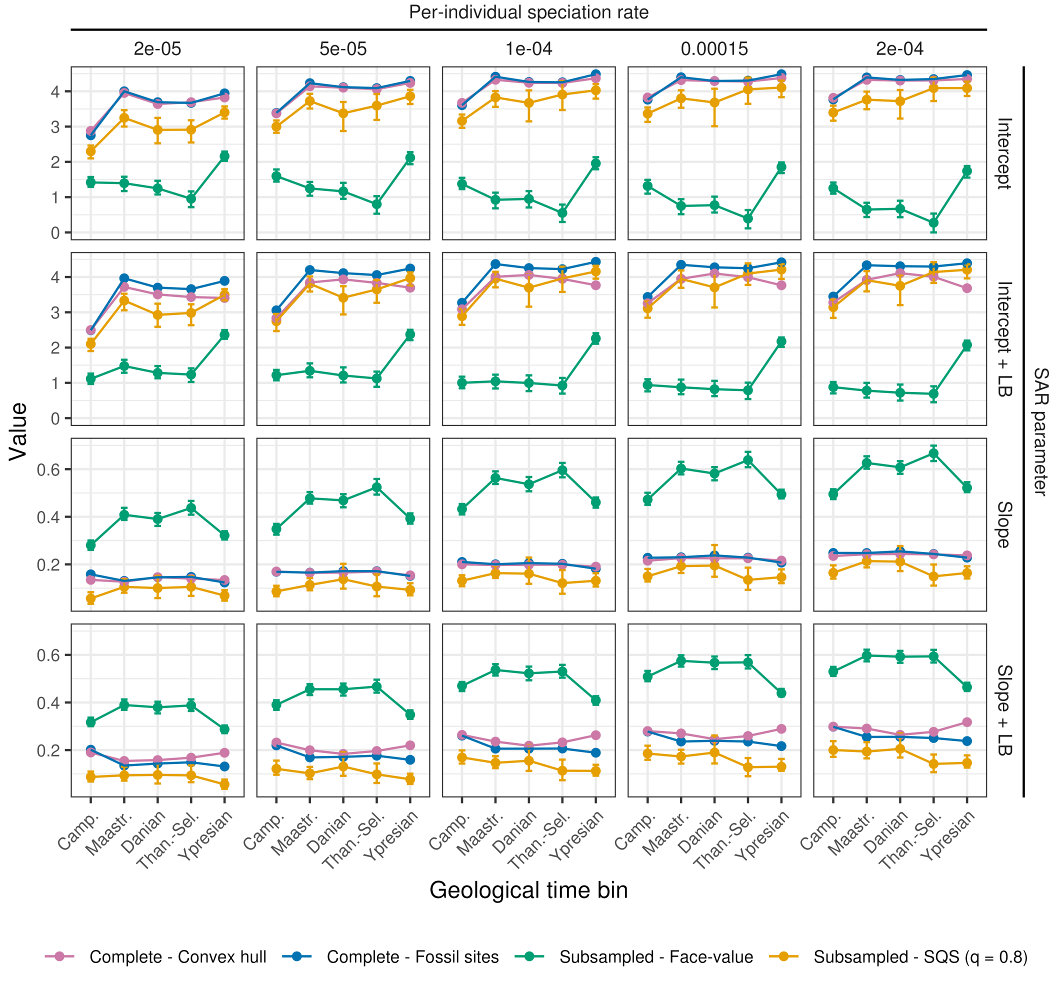


**Fig. S4.** Slopes and intercepts of SENMs for a dispersal distance $\sigma = 50$. OLS regression estimates are shown for SENM simulations that have been subsampled to the region of interest. Columns represent increasing per-individual speciation rates. Rows represent the intercepts and slope estimates of the regressions for the data sets with fossil sites at a great-circle distance of 1000km removed or included (+LB). Distinct colours respectively highlight SAR estimates for the full convex hull with all sites perfectly sampled (purple), complete sampling of all fossil sites (blue), sample sites are subsampled to reflect the number of collections and diversity is measured by face-value counts (green) or SQS (orange). Brackets show the 95% confidence intervals across 100 randomizations.

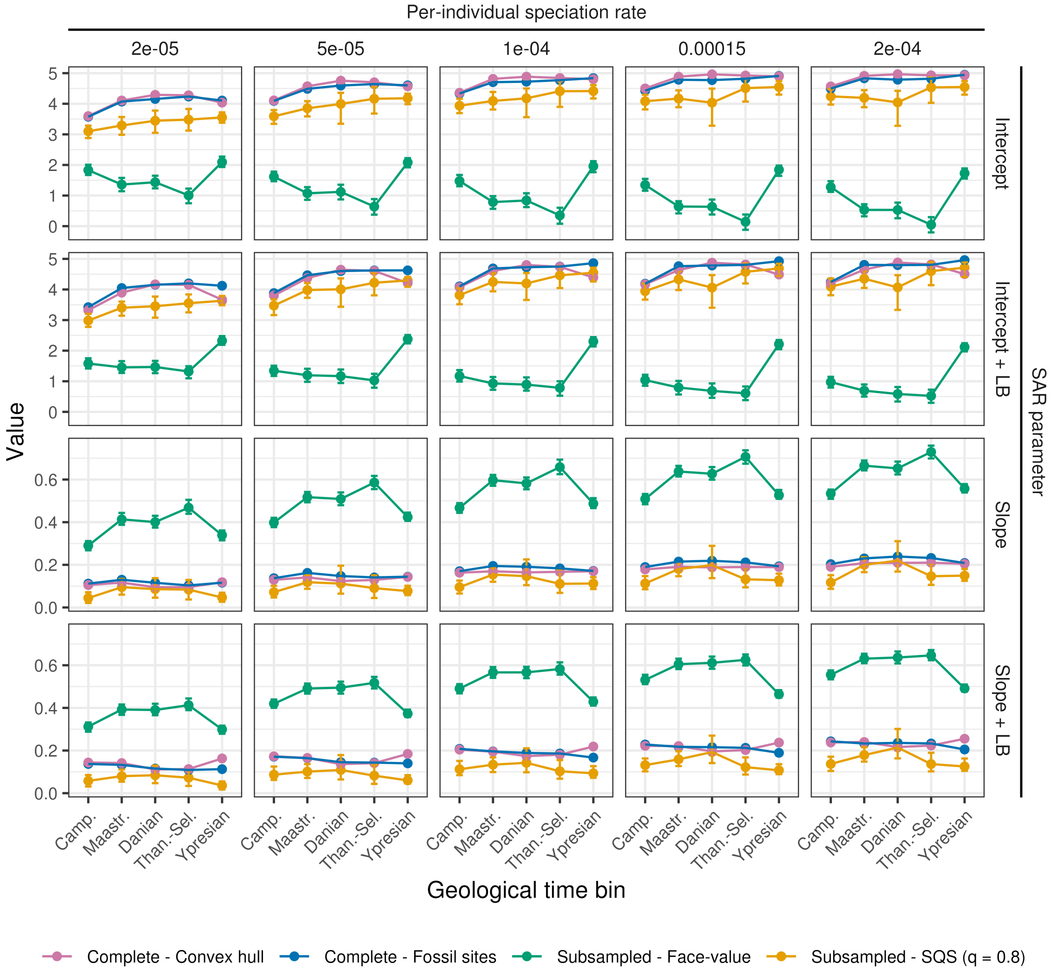


**Fig. S5.** Slopes and intercepts of SENMs for a dispersal distance $\sigma = 100$. OLS regression estimates are shown for SENM simulations that have been subsampled to the region of interest. Columns represent increasing per-individual speciation rates. Rows represent the intercepts and slope estimates of the regressions for the data sets with fossil sites at a great-circle distance of 1000km removed or included (+LB). Distinct colours respectively highlight SAR estimates for the full convex hull with all sites perfectly sampled (purple), complete sampling of all fossil sites (blue), sample sites are subsampled to reflect the number of collections and diversity is measured by face-value counts (green) or SQS (orange). Brackets show the 95% confidence intervals across 100 randomizations.

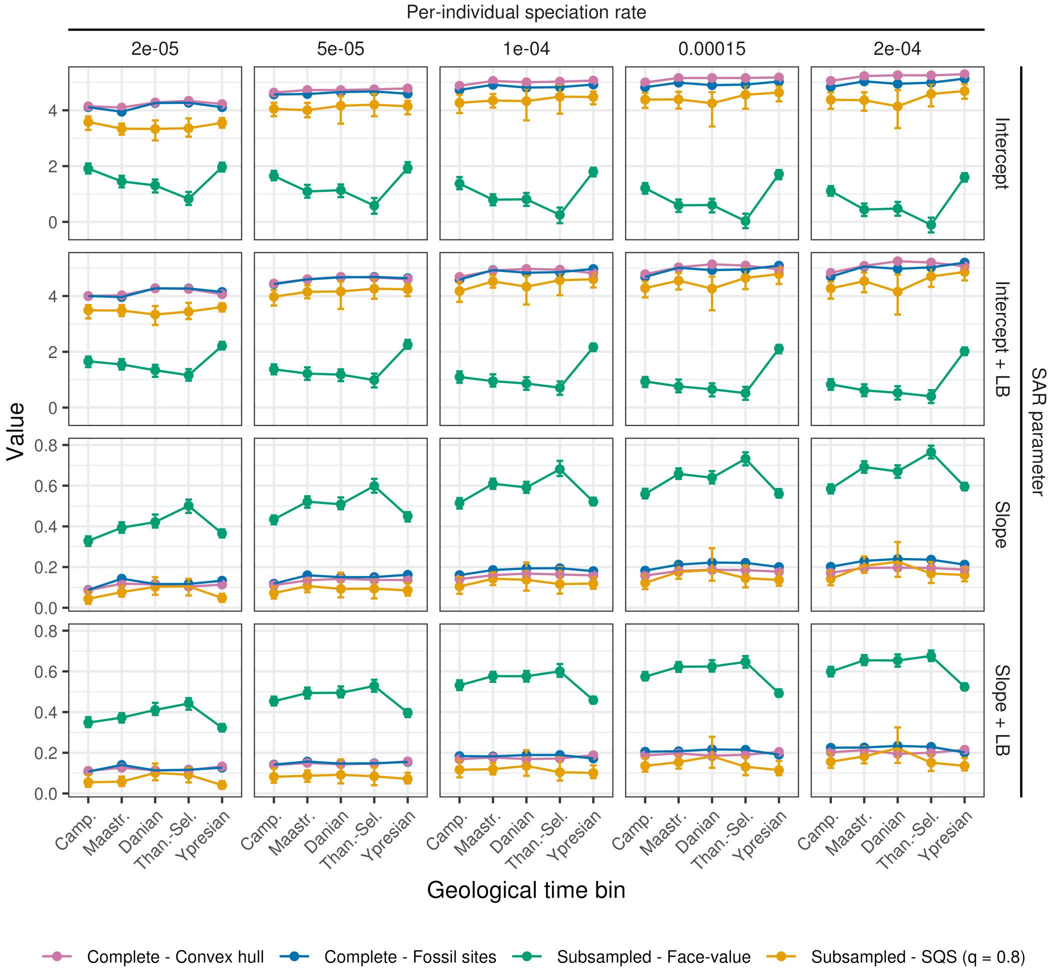


**Fig. S6.** Slopes and intercepts of SENMs for a dispersal distance $\sigma = 150$. OLS regression estimates are shown for SENM simulations that have been subsampled to the region of interest. Columns represent increasing per-individual speciation rates. Rows represent the intercepts and slope estimates of the regressions for the data sets with fossil sites at a great-circle distance of 1000km removed or included (+LB). Distinct colours respectively highlight SAR estimates for the full convex hull with all sites perfectly sampled (purple), complete sampling of all fossil sites (blue), sample sites are subsampled to reflect the number of collections and diversity is measured by face-value counts (green) or SQS (orange). Brackets show the 95% confidence intervals across 100 randomizations.

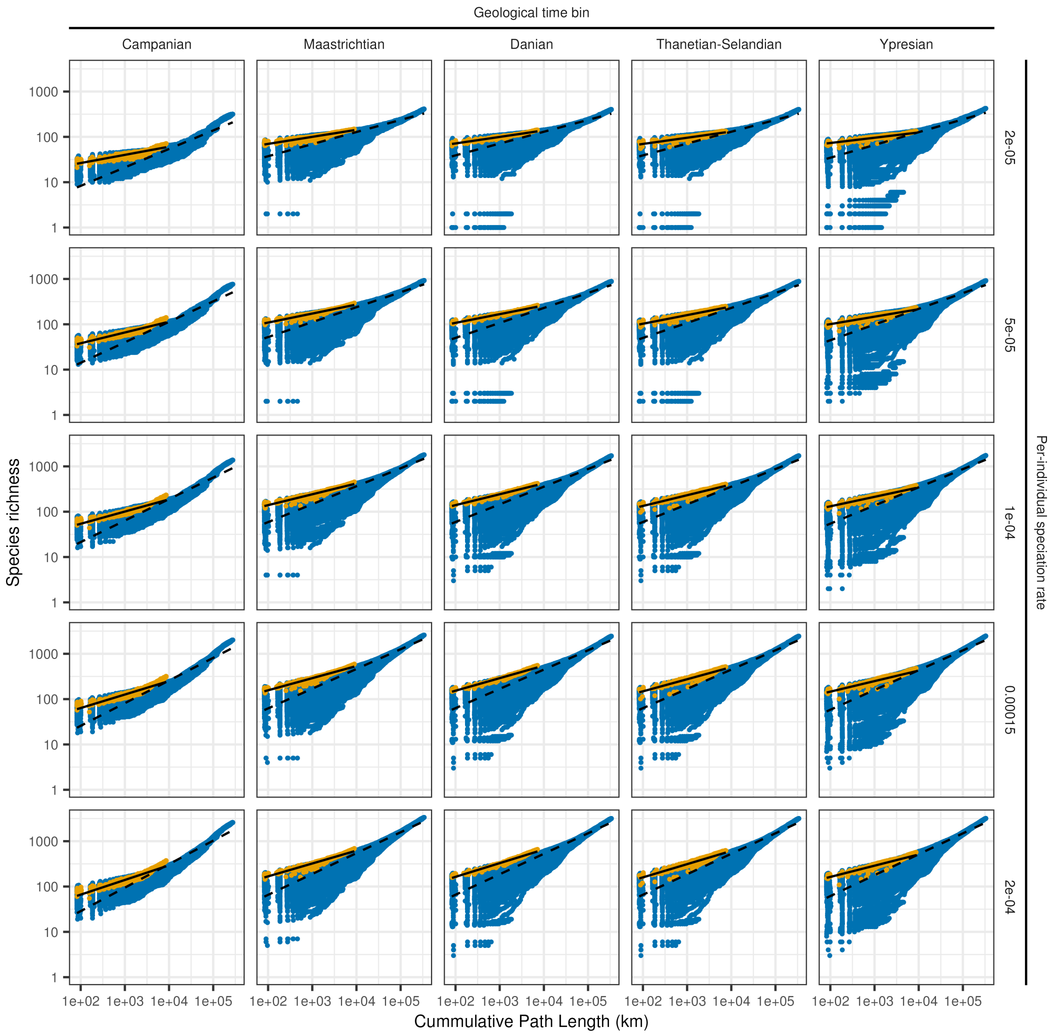


**Fig. S7.** SARs of SENMs for a dispersal distance $\sigma = 25$. Columns and rows respectively highlight the consecutive composite time bins, whereas rows show varying speciation rates. Blue dots show the local-to-regional SAR for the full simulation output across all cells on the North American subcontinent. In contrast, orange dots show the SAR inferred from complete sampling of fossil sites. OLS regression fits are shown on log-log axes for both the full local-to-regional SAR (dashed) and fossil subset (solid).

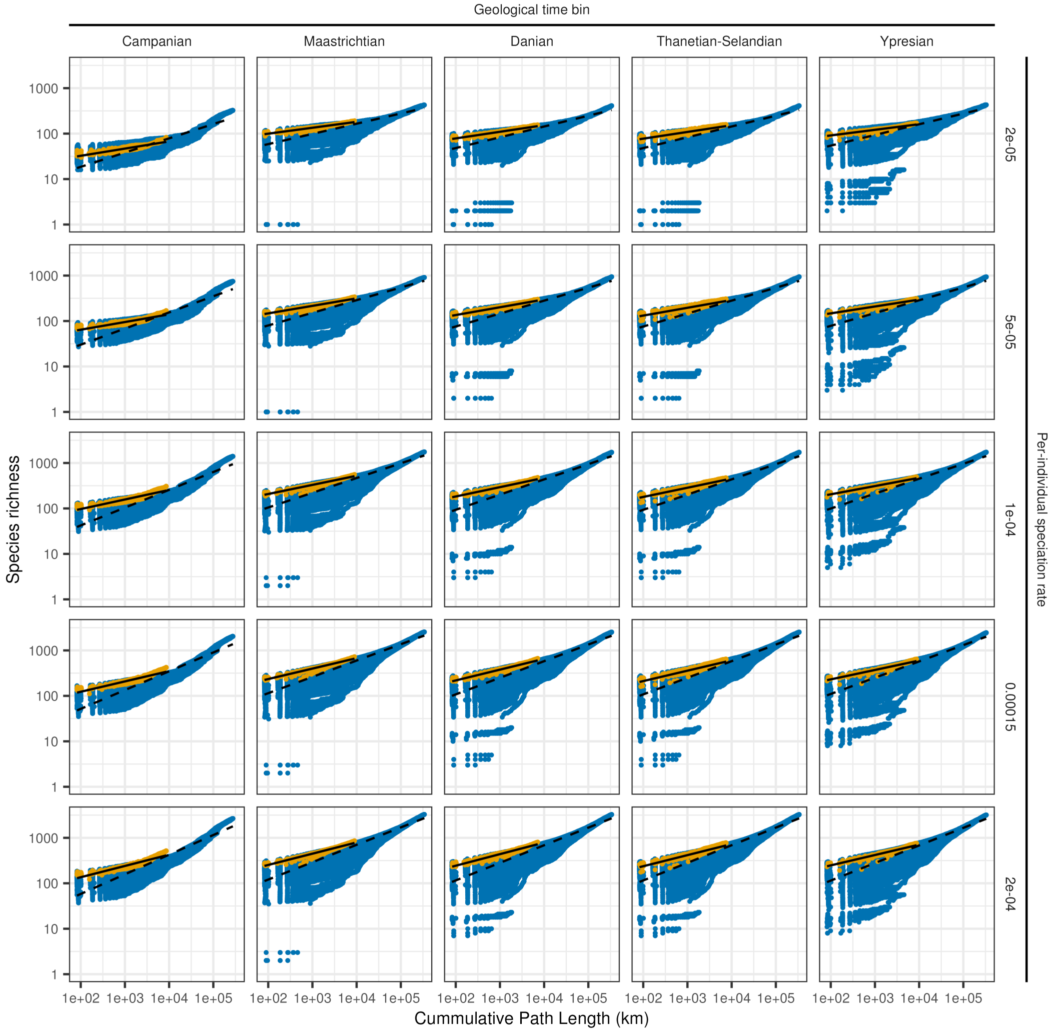


**Fig. S8.** SARs of SENMs for a dispersal distance $\sigma = 50$. Columns and rows respectively highlight the consecutive composite time bins, whereas rows show varying speciation rates. Blue dots show the local-to-regional SAR for the full simulation output across all cells on the North American subcontinent. In contrast, orange dots show the SAR inferred from complete sampling of fossil sites. OLS regression fits are shown on log-log axes for both the full local-to-regional SAR (dashed) and fossil subset (solid).

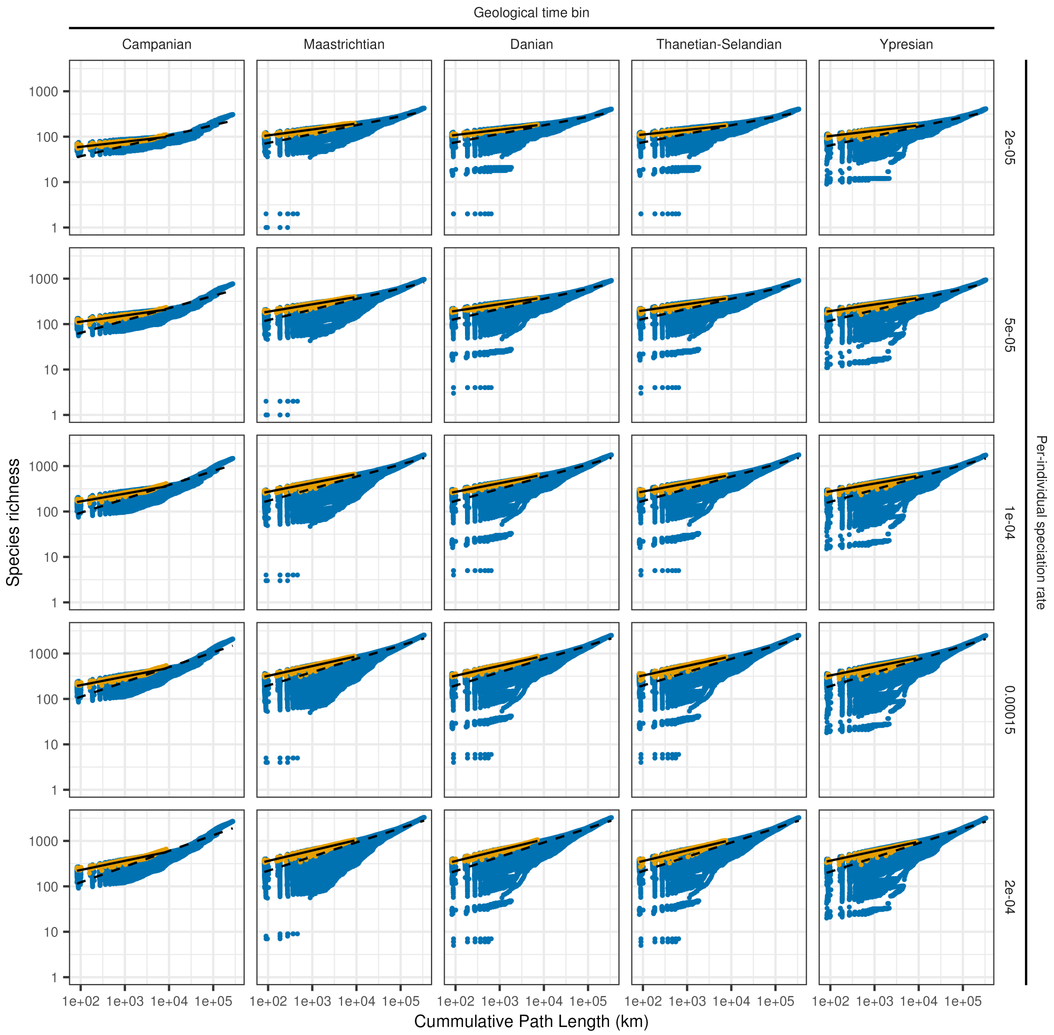


**Fig. S9.** SARs of SENMs for a dispersal distance $\sigma = 100$. Columns and rows respectively highlight the consecutive composite time bins, whereas rows show varying speciation rates. Blue dots show the local-to-regional SAR for the full simulation output across all cells on the North American subcontinent. In contrast, orange dots show the SAR inferred from complete sampling of fossil sites. OLS regression fits are shown on log-log axes for both the full local-to-regional SAR (dashed) and fossil subset (solid).

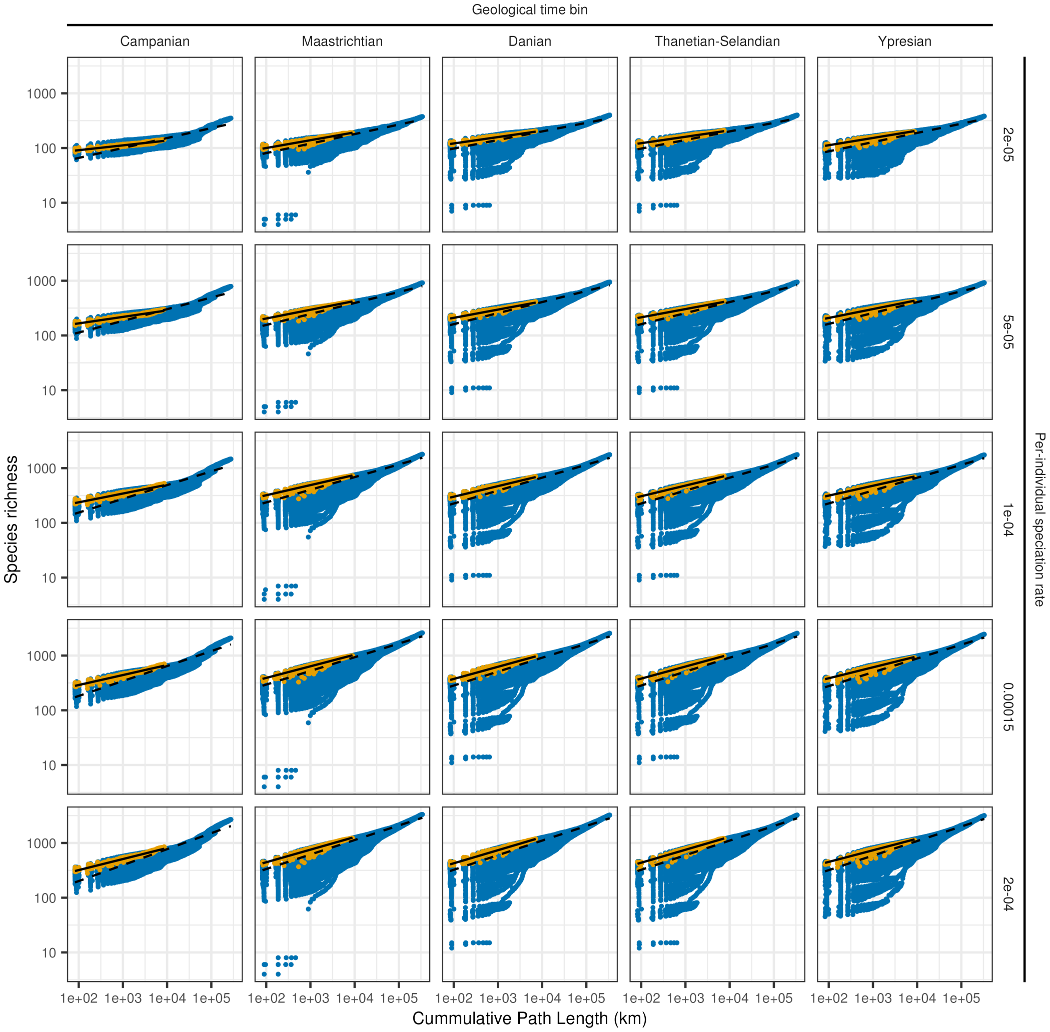


**Fig. S10.** SARs of SENMs for a dispersal distance $\sigma = 150$. Columns and rows respectively highlight the consecutive composite time bins, whereas rows show varying speciation rates. Blue dots show the local-to-regional SAR for the full simulation output across all cells on the North American subcontinent. In contrast, orange dots show the SAR inferred from complete sampling of fossil sites. OLS regression fits are shown on log-log axes for both the full local-to-regional SAR (dashed) and fossil subset (solid).

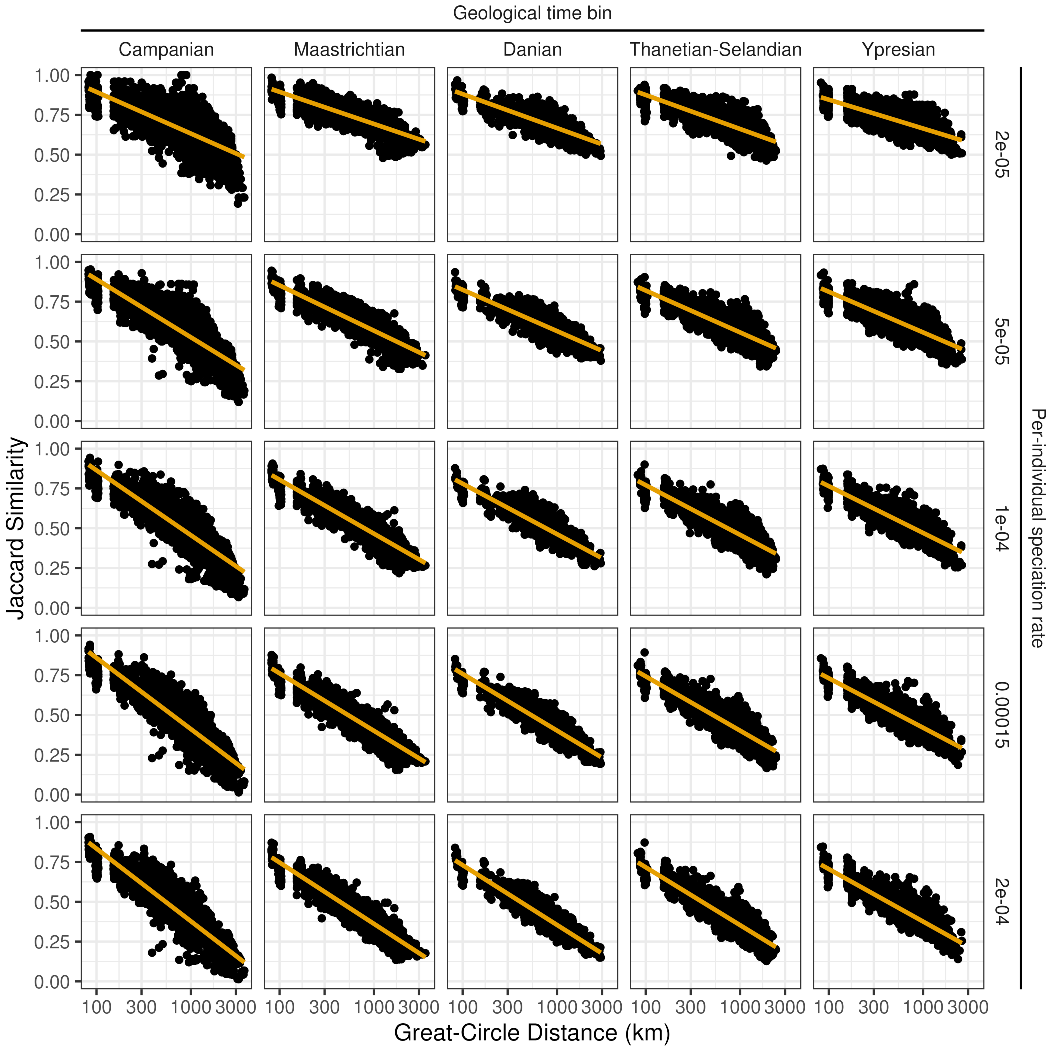


**Fig. S11.** Distance-decay relationships of SENM realisations for dispersal distance $\sigma = 25$. Relationships between Jaccard similarity (i.e. number of shared species divided by the number of unique species in both communities) and great-circle distance are shown for the fossil sites with complete sampling of SENM simulations. Rows represent varying speciation rates, whereas columns show the distinct composite time bins. Orange lines show linear regression fits between similarity and log-transformed geographic distance.

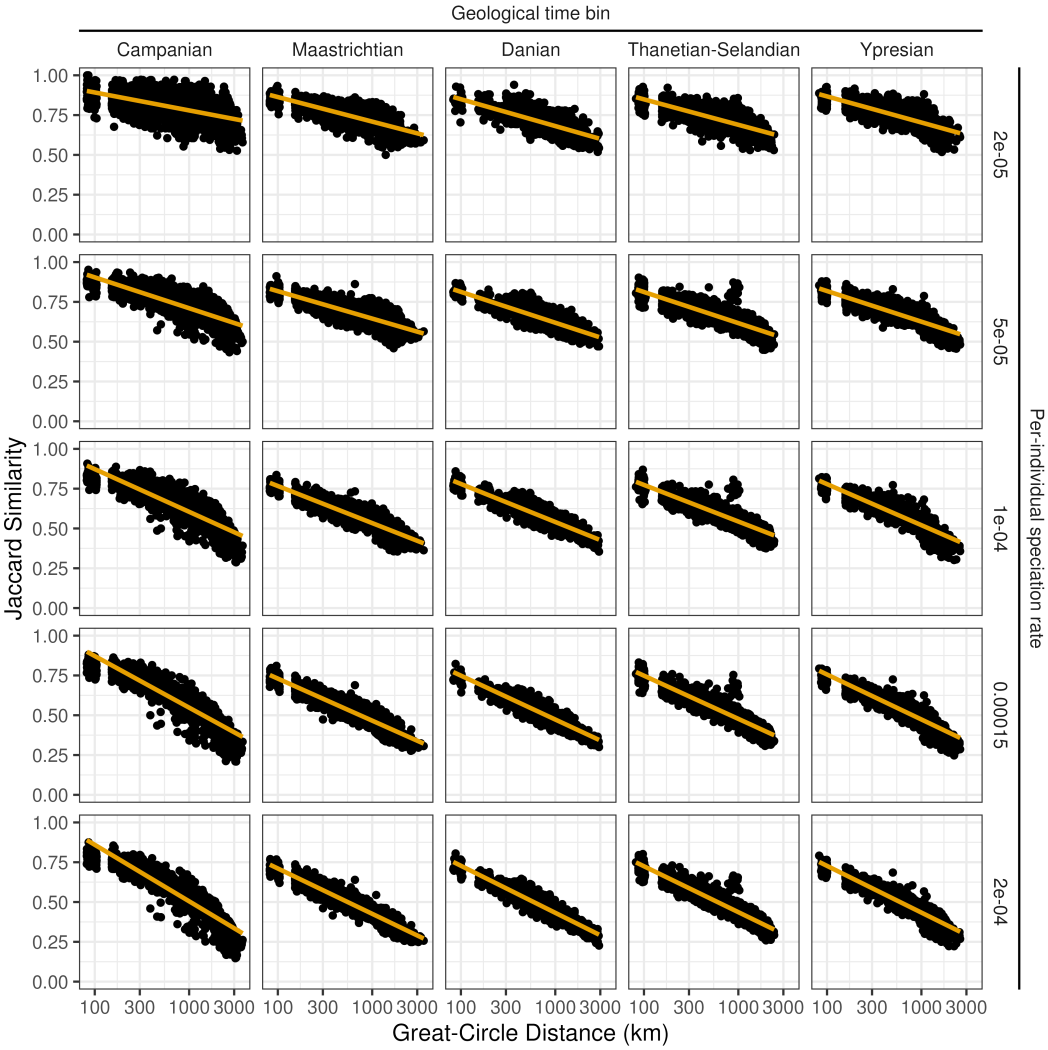


**Fig. S12.** Distance-decay relationships of SENM realisations for dispersal distance $\sigma = 50$. Relationships between Jaccard similarity (i.e. number of shared species divided by the number of unique species in both communities) and great-circle distance are shown for the fossil sites with complete sampling of SENM simulations. Rows represent varying speciation rates, whereas columns show the distinct composite time bins. Orange lines show linear regression fits between similarity and log-transformed geographic distance.

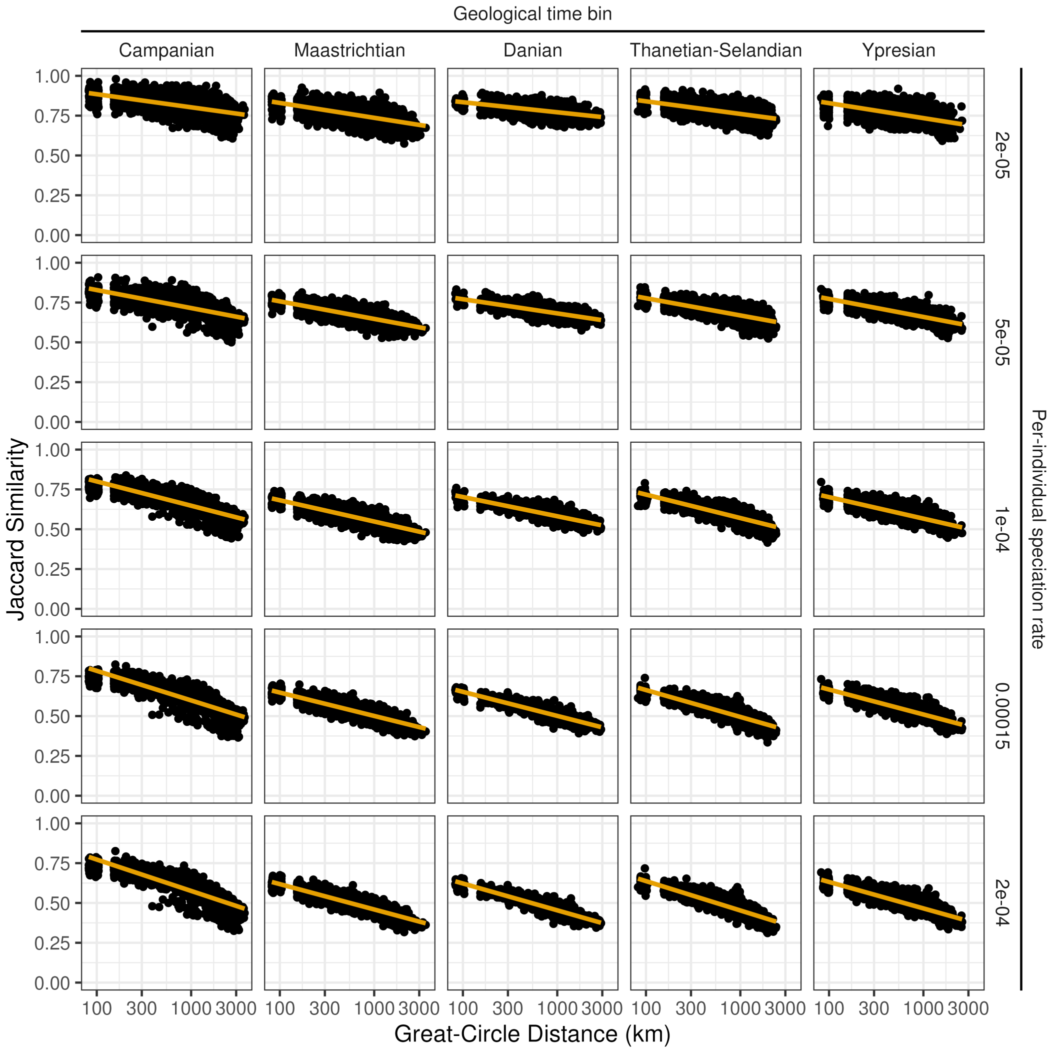


**Fig. S13.** Distance-decay relationships of SENM realisations for dispersal distance $\sigma = 100$. Relationships between Jaccard similarity (i.e. number of shared species divided by the number of unique species in both communities) and great-circle distance are shown for the fossil sites with complete sampling of SENM simulations. Rows represent varying speciation rates, whereas columns show the distinct composite time bins. Orange lines show linear regression fits between similarity and log-transformed geographic distance.

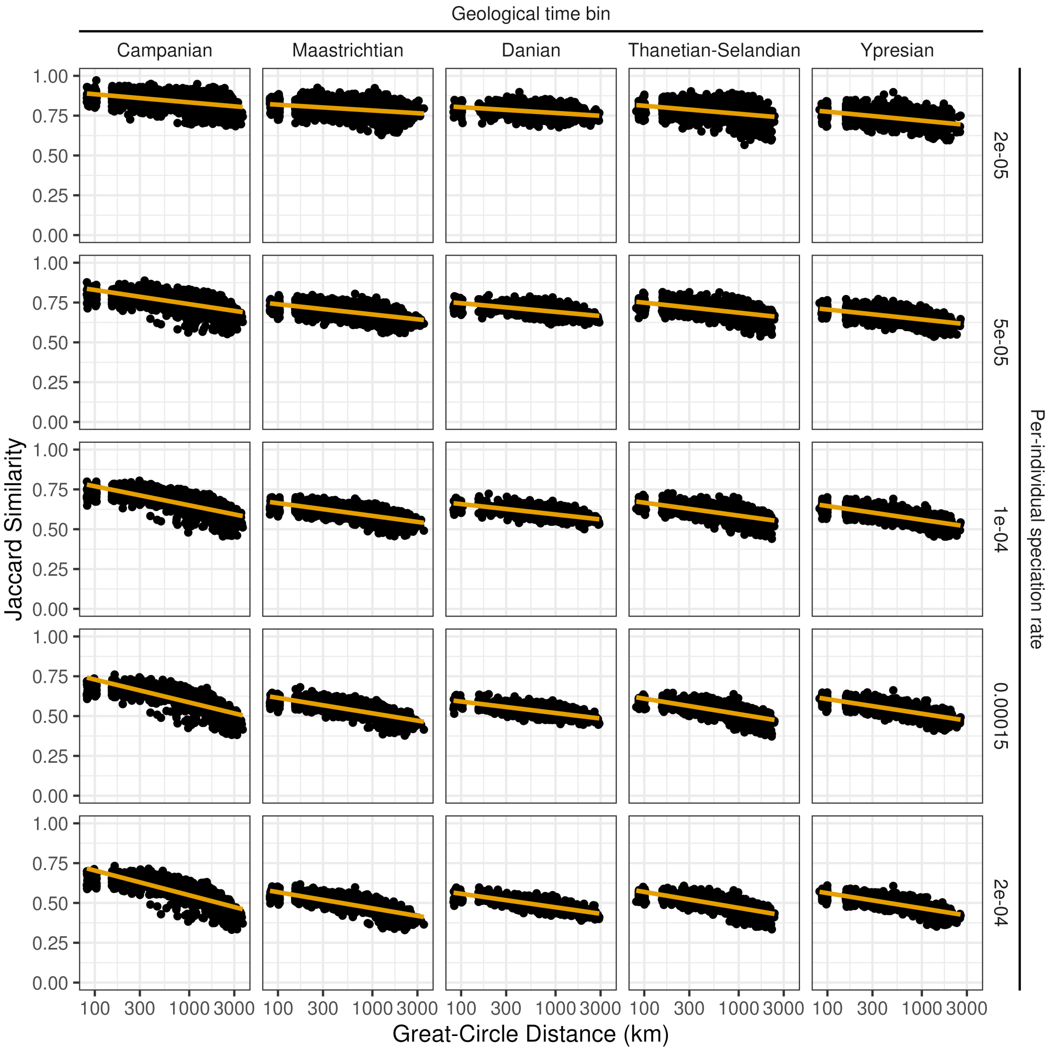


**Fig. S14.** Distance-decay relationships of SENM realisations for dispersal distance $\sigma = 150$. Relationships between Jaccard similarity (i.e. number of shared species divided by the number of unique species in both communities) and great-circle distance are shown for the fossil sites with complete sampling of SENM simulations. Rows represent varying speciation rates, whereas columns show the distinct composite time bins. Orange lines show linear regression fits between similarity and log-transformed geographic distance.

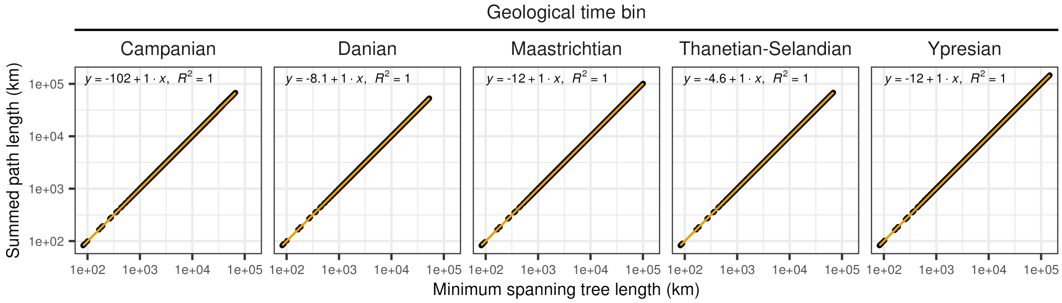


**Fig. S15.** The correlation between minimum spanning tree length (MST) and summed path length. log-transformed distances are shown for the convex hull of all the coordinates per composite time bin. OLS regressions are highlighted by the orange lines. Correlations are exceptionally strong, but MST length is often slightly lower as indicated by the negative intercepts.
